# Region-specific uncoupling of oxygen and glucose metabolism in the human cortex during visual stimulation

**DOI:** 10.1101/2024.10.11.617828

**Authors:** Samira Maria Epp, Antonia Bose, Roman Belenya, Katarzyna Kurcyus, Eric Ceballos Dominguez, Andreas Ranft, Eliana Salas Villa, Moritz Bursche, Igor Yakushev, Christine Preibisch, Gabriel Castrillón, Valentin Riedl

**Affiliations:** Institute of Neuroradiology, Uniklinikum Erlangen, Friedrich-Alexander-University Erlangen-Nuremberg, Erlangen, Germany; Department of Neuroradiology, Neuroimaging Center, Technical University of Munich, University Hospital, Munich, Germany; Graduate School of Systemic Neurosciences, Ludwig-Maximilians-Universität, Munich, Germany; Department of Anesthesiology and Intensive Care Medicine, Technical University of Munich, University Hospital, Munich, Germany; Department of Bioengineering, Universidad de Antioquia, Medellín, Colombia; Department of Nuclear Medicine, Technical University of Munich, University Hospital, Munich, Germany; Research group in Medical Imaging and AI in Healthcare, Biosciences Center, SURA, Medellín, Colombia

## Abstract

The brain primarily relies on oxidative glucose metabolism; however, the oxygen-to-glucose index (OGI) has been suggested to decrease during neuronal activation. Direct evidence for this uncoupling is limited, as the cerebral metabolic rates of oxygen (CMRO₂) and glucose (CMRglc) have not been measured simultaneously in humans under different physiological conditions. We combined multiparametric quantitative BOLD (mqBOLD) imaging with functional PET (fPET) to quantify oxygen and glucose metabolism simultaneously during rest and visual stimulation in a single session. At rest, OGI in the visual cortex was approximately 4.6–5.0, consistent with predominantly oxidative metabolism. During stimulation, CMRglc and cerebral blood flow (CBF) increased, while oxygen extraction fraction (OEF) decreased. CMRO₂ rose only modestly, resulting in a 12% reduction in OGI within the visual cortex. Interestingly, this glycolytic shift extended across the wider cortex (∼8%), but through a different mechanism. While the visual cortex displayed increased CBF and CMRglc with reduced OEF, characteristic of aerobic glycolysis, the remaining cortex showed reduced CBF and CMRO₂ but maintained CMRglc. Our study demonstrates the feasibility of measuring the dynamics of both energy substrates simultaneously in the human brain and reveals a widespread, regionally heterogeneous shift in energy metabolism in response to stimulation. (196)

## Introduction

The human brain stores almost no fuel and depends on the continuous delivery of glucose and oxygen by blood circulation (Dienel, 2019). Most glucose taken up by the brain is oxidized through glycolysis, the tricarboxylic acid cycle and oxidative phosphorylation, while a smaller fraction is converted to lactate after glycolysis and released even when oxygen is abundant, a process termed aerobic glycolysis (Dienel, 2019). Because full oxidation of one glucose molecule consumes six molecules of oxygen, the ratio of the cerebral metabolic rate of oxygen to that of glucose (CMRO₂/CMRglc), the oxygen-to-glucose index (OGI), equals six under stoichiometric conditions (P. Fox et al., 1988). An OGI below six indicates non-oxidative glucose disposal or glycolysis, whereas an OGI above six implies oxidation of alternative substrates such as lactate or ketone bodies (Kersten et al., 1999; Kolb et al., 2021; Pan et al., 2000).

In the resting human cortex, OGI averages approximately 5.5, corresponding to roughly 10% of glucose being metabolized non-oxidatively (Blazey, Snyder, Goyal, et al., 2018). Several PET studies describe a reproducible topography, with the highest non-oxidative fractions in association cortex and the lowest in primary sensory regions (Blazey, Snyder, Su, et al., 2018; Vaishnavi et al., 2010), while others report uniform distributions of glucose oxidation and oxygen extraction across grey matter (Hyder et al., 2016). Independent of its spatial distribution, aerobic glycolysis has been linked to biosynthesis, rapid ATP supply for ion homeostasis, redox buffering and glycogen turnover, particularly in postsynaptic compartments where mitochondria are sparse (Dienel, 2019; Goyal et al., 2014).

The canonical view of activation-induced metabolism derives from early ¹⁵O- and FDG-PET work, in which somatosensory and visual stimulation raised cerebral blood flow (CBF) and CMRglc by 30–50% but CMRO₂ by only ∼5%, implying a steep fall in OGI and a pronounced glycolytic shift (P. Fox et al., 1988; P. T. Fox & Raichle, 1986). This finding has been influential yet also disputed: ³¹P-MRS in humans indicated ATP turnover during activation far in excess of what non-oxidative pathways could supply (Rango et al., 1997), and ¹³C-NMR during intense neuronal activation in rodents showed glucose oxidation tightly coupled to glutamatergic flux (Patel et al., 2004). ¹⁷O-MRS in cats reported CMRO₂ increases above 30% within activated visual cortex, accompanied by CMRO₂ decreases in surrounding tissue, and highlighted the dependence of relative responses on baseline metabolic state (Zhu et al., 2009). Calibrated fMRI has variously supported near-linear CBF-CMRO₂ coupling without a plateau (Hoge et al., 1999) and an upper bound of ∼30% for ΔCMRO₂ with CBF increases exceeding it (W. Chen et al., 2001). Intermediate CMRO₂ estimates have been reported with dual-tracer PET (Blomqvist et al., 1994), quantitative fMRI under graded hypoxia (Tuunanen et al., 2006), and a range of MRI-based CMRO₂ models (A. Lin et al., 2008; A.-L. Lin et al., 2010, 2012; Qin et al., 2011; Zhu et al., 2018). Biophysical and metabolic modeling has likewise been used both to reconcile and to challenge a large flow– oxidation mismatch (Calvetti et al., 2018; Gustafsson et al., 2024; Lundengård et al., 2016; Mintun et al., 2001). Thus, the magnitude of activation-induced uncoupling remains unresolved (P. T. Fox, 2012; Hyder et al., 2010; Mangia et al., 2009).

A substantial part of this disagreement is methodological rather than physiological. Almost all prior estimates of activation-induced OGI combine CMRO₂ and CMRglc acquired in separate sessions, from separate cohorts, or in anaesthetised animals (P. Fox et al., 1988; P. T. Fox & Raichle, 1986; Hyder et al., 2016; Leenders et al., 1990; Madsen et al., 1999; Vafaee et al., 2012). Because CMRO₂, CMRglc and CBF all vary with arousal, glycaemia, anaesthesia and baseline metabolic state, ratios formed across acquisitions inherit this variance, and no design has so far permitted OGI to be tracked across physiological states within the same subjects.

Hybrid PET/MR, together with dynamic sampling of PET- and MR-data, makes this feasible. Functional PET (fPET) with a bolus-plus-constant-infusion of [¹⁸F]FDG resolves task-related changes in CMRglc within a single scan (Hahn et al., 2016; Jamadar et al., 2019; Rischka et al., 2018; Villien et al., 2014), while multiparametric quantitative BOLD (mqBOLD) derives voxel-wise oxygen extraction fraction (OEF) and CMRO₂ from T2, T2*, cerebral blood volume (CBV) and arterial spin labelling perfusion (Christen et al., 2012; Kaczmarz et al., 2020), avoiding both gas challenges (Blockley et al., 2013a; Kim et al., 1999) and short-lived ¹⁵O tracers (Herscovitch et al., 1983; Mintun et al., 1984). We have previously applied this framework to map the energetic architecture of the resting human connectome (Castrillon et al., 2023) and to show that BOLD signal changes can oppose the direction of oxygen metabolism across the cortex (Epp et al., 2026).

Here we combined fPET and mqBOLD in the same session and the same participants to quantify CMRglc and CMRO₂ simultaneously during alternating blocks of full-field visual stimulation and fixation. We asked three questions: (i) whether resting OGI in visual and extra-visual cortex is consistent with published whole-brain estimates; (ii) how large the stimulation-induced OGI decrease is when oxygen and glucose metabolism are measured concurrently rather than across sessions; and (iii) whether any such shift is confined to task-engaged regions or extends across the cortex. By removing the between-session variance that has constrained earlier work, this design provides a direct test of how far functional activation shifts the human cortex away from fully oxidative glucose metabolism.

## Results

In this study, we combined simultaneous FDG-fPET and mqBOLD MRI to quantify CMRglc and CMRO₂ metabolism during a visual stimulation paradigm with alternating ∼6-min blocks of moving checkerboard stimulation (STIM) and visual fixation (REST) (Fig. 1A). The multimodal design allowed us to capture complementary aspects of neurovascular and neurometabolic coupling: BOLD-fMRI served as a functional localizer of task-evoked signal changes that primarily reflect changes in deoxyhemoglobin driven by the interplay of CBF, CBV, and oxygen consumption (Fig. 1B). mqBOLD combined T2, T2*, dynamic susceptibility contrast (DSC) CBV, and pseudo-continuous arterial spin labeling (pCASL) CBF measurements to estimate OEF and derive voxelwise CMRO₂ as a quantitative proxy of oxidative metabolism (Fig. 1C). FDG-fPET and arterial sampling provided a dynamic estimate of CMRglc, reflecting regional glucose uptake and utilization independently of blood oxygenation (Fig. 1D). Together, these measures enabled us to compare where visual stimulation altered hemodynamics, oxidative metabolism, and glucose metabolism compared to visual fixation, and how these responses together shape the OGI. All baseline values (at REST) of the quantitative imaging parameters were within physiologically plausible ranges (Table 1).

**Figure 1:**
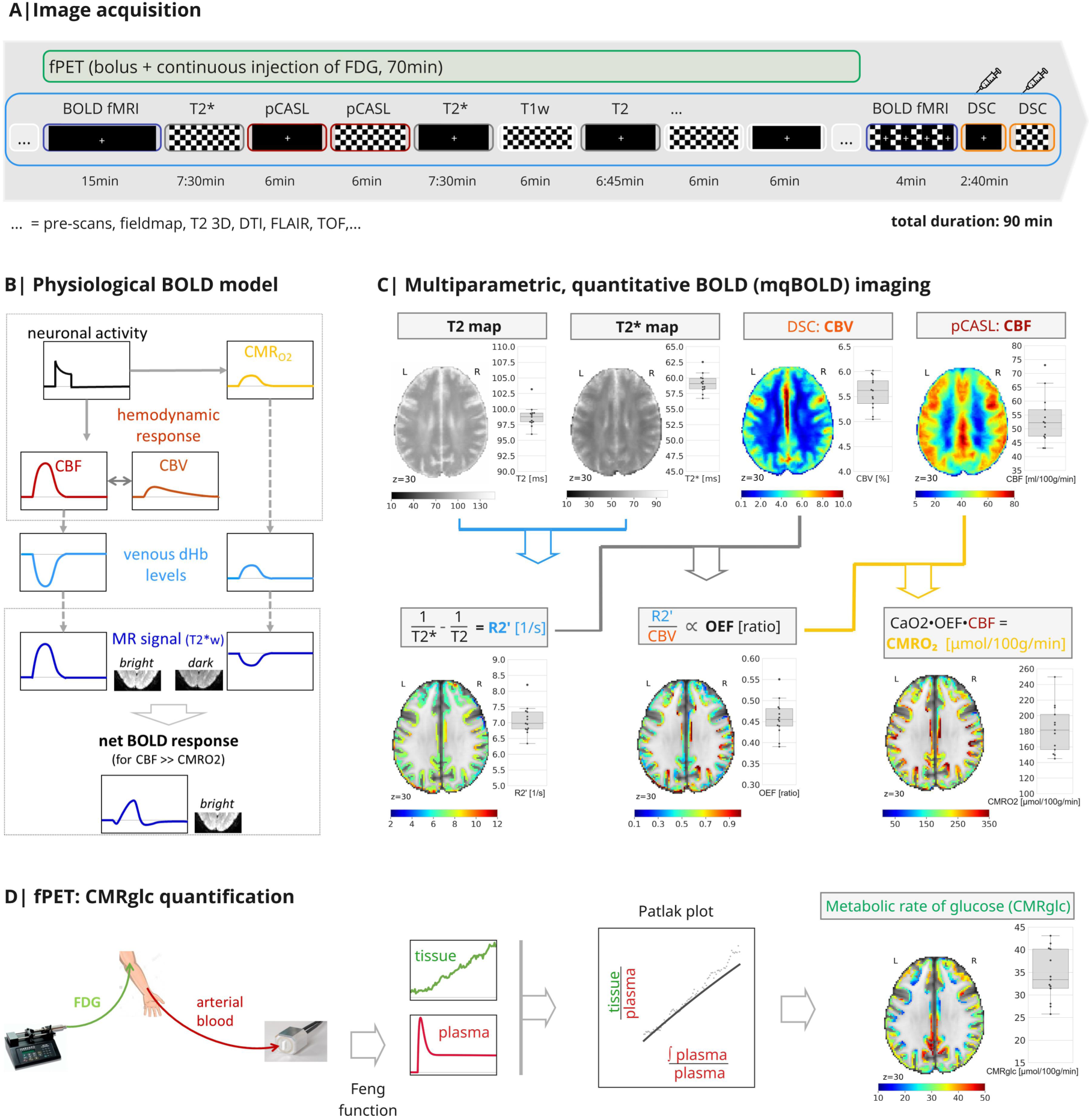
Data acquisition and physiological framework. **A**) Simultaneous fPET (green) and MRI (blue) were acquired in healthy human subjects over 90 min (70min for fPET) during alternating checkerboard and fixation blocks. [¹⁸F]FDG was administered using a bolus-plus-infusion protocol (20% bolus, 80% continuous infusion). BOLD-fMRI and dynamic susceptibility contrast (DSC) scans, the latter following gadolinium administration, were acquired at the end of the session. **B**) Neuronal activation increases CMRO₂, which raises venous deoxyhemoglobin (dHb), while in parallel increasing CBF, which reduces dHb. Because the relative increase in CBF exceeds that of CMRO₂, the net dHb concentration decreases, producing a positive BOLD signal response. **C**) Multiparametric quantitative BOLD (mqBOLD) estimates CMRO₂ from T2, T2*, cerebral blood volume (CBV) from DSC, and cerebral blood flow (CBF) from pCASL, generating voxelwise maps of the cerebral metabolic rate of oxygen (CMRO₂). CaO2 = arterial oxygen content. Axial brain slices show subject-average, non-thresholded parameter maps during REST, and boxplots show individual-subject median gray-matter values. **D**) fPET quantifies condition-specific CMRglc from dynamic PET data using an arterial input function, a two*-* compartment kinetic model, and Patlak analysis.

**Table 1:** Baseline (REST) values, median (IQR) across subjects, within GM.

| R2' | CBV | OEF | CBF | CMRO<sub>2</sub> | CMRglc |
| --- | --- | --- | --- | --- | --- |
| [1/s] | [%] | [ratio] | [ml/100g/min] | [μmol/100g/min] | [μmol/100g/min] |
| 7.0 | 5.6 | 0.46 | 52.2 | 181.5 | 33.4 |
| (0.5) | (0.5) | (0.04) | (9.6) | (44.9) | (8.7) |

Based on canonical neurovascular coupling, we expected visual stimulation to increase CBF and decrease OEF, because the rise in blood flow exceeds the increase in oxygen consumption, thereby producing a positive BOLD response. We further expected increases in both CMRO₂ and CMRglc, with a larger relative increase in CMRglc, consistent with a task-evoked reduction in OGI.

### The BOLD response reflects spatially heterogeneous neurovascular and neurometabolic processes

We identified task-responsive regions by comparing voxel-wise maps of BOLD, CMRglc, CMRO₂, CBF, and OEF between REST and STIM and found significant responses across all parameter maps (p < 0.05, corrected; Fig. 2A). The spatial extent of task-evoked effects varied across modalities. CMRO₂ and CBF responses were relatively widespread throughout the cortex, whereas BOLD and CMRglc changes were concentrated in the visual cortex. OEF was also primarily located in the visual cortex. Within the visual cortex, all parameters except OEF increased, whereas outside the visual cortex, CMRO₂ and CBF also decreased. Thus, visual stimulation elicited broad hemodynamic adjustments but more localized positive metabolic responses.

**Figure 2:**
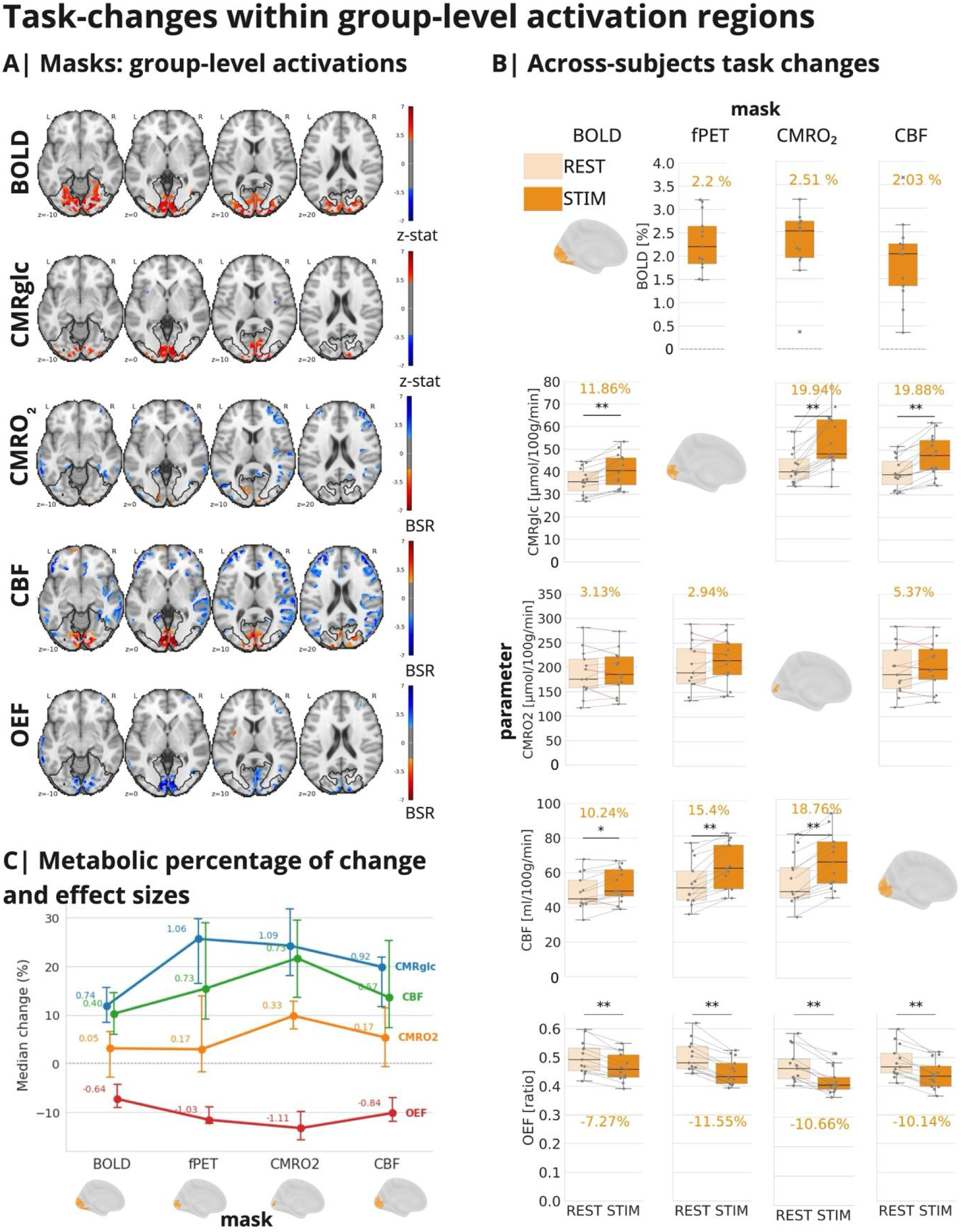
Task-Evoked Responses During Visual Stimulation. **A**) Group-level maps illustrate the contrast between the STIM and REST conditions, based on second-level GLM analyses for BOLD and CMRglc, and the first PLS latent variable for CMRO₂, CBF, and OEF (N=13 for BOLD, CBF, OEF, and CMRO₂; N=15 for CMRglc). Warm colors indicate increased parameter values during STIM compared to REST, while cool colors indicate the opposite (GLM: Z > 3.2, controlling for voxel-wise FDR of p < 0.05; PLS: |BSR| > 2, p < 0.05, 2000 permutations). Black contours outline the visual cortex. Within the visual cortex, BOLD, CBF, CMRO₂, and CMRglc increased, while OEF decreased. **B**) Hemodynamic and metabolic changes across subjects within parameter-specific masks. All parameters showed consistent effects, except for CMRO₂, which did not exhibit significant changes across subjects. **C**) Summary: The group-median percent changes within masks displayed strongest changes in CMRglc, followed by CBF, OEF and CMRO₂ across masks. Error bars show the 95% bootstrap confidence interval, and the numbers indicate Hedges’ g effect size.

### Individual task-evoked changes in the visual cortex

We next tested whether the group-level findings were also evident at the subject level by comparing median parameter values between REST and STIM within task-evoked visual regions of interest (ROIs). Consistent across parameter-specific ROIs, the most robust changes were observed in CMRglc, CBF, and OEF, whereas increases in CMRO₂ were not significant (Fig. 2B,C). Specifically, within the BOLD-defined mask (Fig. 2B, first column), CMRglc increased by 4.4 μmol/100 g/min (11.9%), CBF increased by 4.6 ml/100 g/min (10.2%), and OEF decreased by 0.04 (−7.3%), whereas CMRO₂ increased by 6.2 μmol/100 g/min (3.1%) but did not reach significance (p = 0.6). Results were similar for masks derived from the other parameter maps (Fig. 2B, columns 2-4). Results also remained consistent when including only the 11 subjects with DSC data available from both conditions (Fig. S1). Furthermore, to assess the impact of CBV assumptions on CMRO₂ estimates, we recalculated group-level CMRO₂ responses without correcting for task-evoked CBV changes. This yielded substantially larger increases in CMRO₂ (7-17% across ROIs). Importantly, OGI results remained unchanged, showing significant reductions in all task-related ROIs and throughout the remaining cortex (Fig. S2), indicating that the main conclusions are robust to larger CMRO₂ estimates when assuming no task-evoked CBV changes. Taken together, the largest task-evoked percentage changes were observed in CMRglc (12-26%) and CBF (10-22%), followed by OEF (−7 to −13%) and CMRO₂ (3-10%) (Fig. 2C).

### Task-evoked glucose metabolic response is predominantly glycolytic

To characterize the balance between oxygen and glucose metabolism, we analyzed changes in the OGI, defined as CMRO₂/CMRglc. An OGI of approximately 6 reflects fully oxidative glucose metabolism, whereas values below 6 indicate a relative shift toward non-oxidative, glycolytic metabolism. In the metabolic ROIs, the mean OGI at rest was around 5.0 (95% CI: 4.5-5.8) within the CMRglc mask and 4.6 (95% CI: 4.1-5.2) within the CMRO₂ mask. Visual stimulation further shifted metabolism toward glycolysis within these visual ROIs. OGI decreased by 12.8% (95% CI: 7.3-18.8%) within the CMRglc mask and by 12.4% (95% CI: 8.4-17.3%) within the CMRO₂ mask. Similarly, mean OGI at rest was around 5.0 (95% CI: 4.5-5.7) within the BOLD mask and decreased by 7.9% (95% CI: 5.0-14.1%) after visual stimulation (Fig. 3A). These reductions indicate that glucose utilization increased more strongly than oxygen consumption in task-engaged visual cortex.

**Figure 3:**
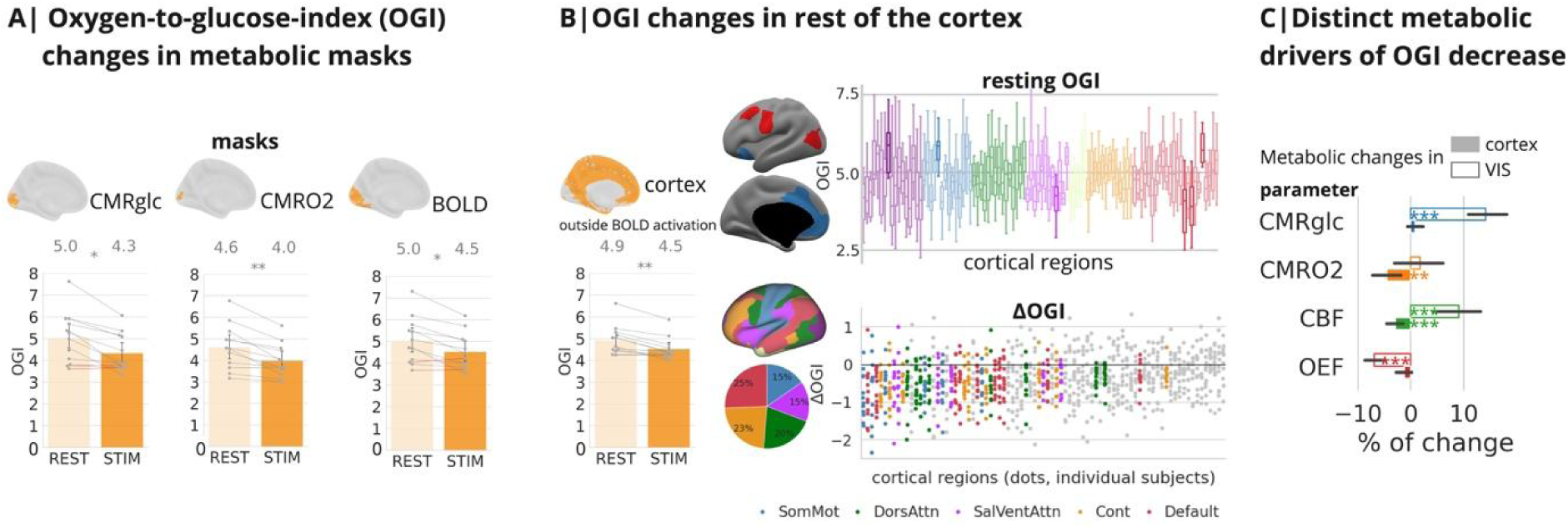
Oxygen-to-glucose index (OGI) during visual stimulation. **A**) OGI decreased significantly during STIM relative to REST within both task-evoked metabolic ROIs (CMRglc mask, left; CMRO₂ mask, middle; BOLD mask, right; p < *0.05, **0.01), indicating a shift toward glycolytic metabolism. **B**) Left: OGI reductions extended beyond the visual cortex (left bar plots; **p < 0.001). Upper-right: resting OGI did not significantly differ across the whole cortex (p = 0.43). Only six regions had an OGI significantly different from the mean resting value, as shown on the brain surface. Red shows OGI above the mean, while blue indicates below the mean. These regions are also highlighted in the boxplot, colored by their functional network assignment. Lower-right: at the regional level, 47% of cortical ROIs showed significant OGI decreases (p<0.05, ΔOGI≠0, FDR-corrected; strip plot), with ROIs colored by their functional network assignment (surface map); the distribution of significant effects across networks is summarized in the pie chart (bottom-center). **C**) Distinct metabolic mechanisms underlie the OGI decrease: in the visual cortex, increased CBF and CMRglc, coupled with reduced OEF, drive a pronounced shift toward glycolysis, whereas the rest of the cortex exhibits a weaker glycolytic shift, driven by reduced CBF and CMRO₂, despite stable CMRglc and OEF (right bar plots; p < **0.01, ***0.0001). Error bars in bar plots represent the 95% bootstrap confidence interval. Network abbreviations: Cont = control, Default = default mode, DorsAttn = dorsal attention, SalVenAttn = salience ventral attention, SomMot = somatomotor, VIS = visual.

Given the widespread stimulation-related changes in CBF and CMRO₂ beyond the visual cortex, we extended the OGI analysis to include the rest of the cortex. Resting OGI outside the visual cortex had a mean of 4.9 (95% CI: 4.6-5.4; Fig. 3B, left) and did not significantly vary across major functional networks (repeated-measures ANOVA: F(6, 66) = 1.01, *p* = 0.43, ηg² = 0.01; Fig. 3B, upper-right). Region-wise OGI showed substantial subject-wise variation (standard deviation: mean = 0.85, 95% CI: 0.81-0.89), yet three frontal regions showed significantly reduced OGI, and one frontal and two sensory cortex regions exhibited significantly increased OGI during REST compared with the average cortex (p<0.05, OGI≠4.9 one-sample t-test, FDR-corrected; Fig. 3B, upper-right). During stimulation, OGI decreased by 8% (95% CI: 4.8-10.4%) beyond the visual cortex (Fig. 3B, left). This effect was spatially distributed rather than driven by occipital regions. Specifically, 47% of cortical ROIs showed significant (p<0.05, ΔOGI≠0 one-sample t-test, FDR-corrected) condition-related OGI changes, distributed similarly across major functional brain networks (Fig. 3B, lower-right).

Together, these findings reveal a widespread stimulus-induced shift toward glycolytic metabolism, particularly in the visual cortex but also throughout the broader cortex. However, the underlying metabolic profiles in the visual cortex differ from those in the rest of the cortex (Fig. 3C). In the visual cortex (BOLD mask), aligned with aerobic glycolysis, both CMRglc and CBF increased, while OEF decreased (p<0.0001; ΔOGI≠0 one-sample t-test; FDR-corrected). This indicates that glucose supply and uptake increased more strongly than oxygen utilization. In contrast, the rest of the cortex exhibited concurrent reductions in CBF and CMRO₂ (p<0.01; ΔOGI≠0; one-sample t-test; FDR-corrected), with no significant changes in CMRglc and OEF. This suggests a reduction in oxidative metabolism despite preserved glucose utilization.

## Discussion

By quantifying oxygen and glucose metabolism within a single imaging session, this study provides direct in vivo evidence that task-evoked activity shifts the cortical oxygen-to-glucose balance toward glycolysis. In the visual cortex, stimulation produced the expected neurovascular response, increased CBF and reduced OEF, together with a disproportionate rise in CMRglc relative to CMRO₂ and a corresponding fall in OGI. A smaller but consistent OGI reduction extended across the wider cortex. Critically, this cortex-wide shift did not reflect a single mechanism: whereas task-engaged visual cortex showed genuine glycolytic upregulation, the remaining cortex reached a lower OGI through reduced oxidative metabolism with preserved glucose use. A shift toward glycolysis is therefore a general accompaniment of functional activation, but it emerges from distinct combinations of flow, oxygen, and glucose metabolism across the cortex.

The dissociation between widespread hemodynamic responses and more localized metabolic changes underscores the complex interpretation of the BOLD signal (Ekstrom, 2021; Mark et al., 2015). Increases in CMRglc and CMRO₂ were largely confined to task-engaged regions, indicating that metabolic demand tracks local neuronal activity (Herman et al., 2025; Wolfe et al., 2024; Yu et al., 2018). Outside the visual cortex, by contrast, widespread CBF reductions were accompanied by decreases in CMRO₂ consistent with a supply-limited framework of brain metabolism (Herculano-Houzel & Rothman, 2022) and with prior reports of activation-related CBF redistribution and reduced oxygen metabolism (Bruckmaier et al., 2020; Ghatan et al., 1998; Kastner & Ungerleider, 2000). Together, these observations indicate that the BOLD response reflects a combination of localized neurometabolic activity and broader vascular dynamics rather than a direct readout of regional energy consumption, a distinction we have recently shown can even reverse the apparent sign of oxygen metabolism (Epp et al., 2026).

In the visual cortex, the concurrence of increased CMRglc and CBF with decreased OEF indicates that activation draws more on glucose supply and glycolytic processing than on oxidative metabolism. This pattern is consistent with earlier [¹⁸F]FDG- and [¹⁵O]-PET work showing that glucose use and blood flow rise more steeply than oxygen consumption during sensory and cognitive tasks (P. Fox et al., 1988; P. T. Fox & Raichle, 1986; Vlassenko et al., 2018), the hallmark of activation-induced aerobic glycolysis (DiNuzzo et al., 2024; Herculano-Houzel & Rothman, 2022). The absence of significant CMRO₂ changes in ROIs defined by parameters other than CMRO₂ suggests that oxidative responses remain spatially restricted to the most strongly engaged tissue, whereas CMRglc, CBF, and OEF changes extend further. Accordingly, CMRglc, CBF, and OEF proved to be the most sensitive indicators of neurovascular–neurometabolic coupling during visual stimulation. Notably, OEF may be particularly attractive for future studies, as it can be estimated without contrast agents using quantitative susceptibility mapping–based approaches (Zhang et al., 2015), which have already been successfully evaluated in clinical populations (Zhuang et al., 2026).

The magnitude of task-evoked changes was broadly consistent with the literature, with the notable exception of CMRO₂. Across activation ROIs, CMRglc increased by 12–26% and CBF by 10–22%, whereas CMRO₂ rose only modestly (≈3–10%) and did not reach significance except within its own defining ROI. This small oxidative response sits at the lower bound of the wide range reported previously — from ≈5% in early PET (P. Fox et al., 1988; P. T. Fox & Raichle, 1986) to ≈11–30% in later calibrated-fMRI and PET studies (Davis et al., 1998; Donahue et al., 2009; Kim et al., 1999) — which were typically accompanied by much larger CBF increases (30–63%). Because those higher estimates were often derived from smaller, activation-centered ROIs that tend to amplify effect sizes, part of the discrepancy is likely methodological; the modest CBF changes measured here, together with a strong increase in total CBV (see Limitations) further constrain our ΔCMRO₂ estimates. Nevertheless, OGI reductions remained significant despite substantially larger CMRO₂ estimates when assuming no task-evoked CBV changes. Yet, the relatively greater rise in CBF than CMRO₂ nonetheless reproduces the OEF decrease that underlies the positive BOLD response (Buxton, 2010; Leithner & Royl, 2014), and the CMRglc increase confirms glucose metabolism as a sensitive marker of visual activation (Hahn et al., 2016; Newberg et al., 2005; Rischka et al., 2018; Villien et al., 2014).

At rest, OGI in the visual cortex (4.8–5.0) was somewhat below the whole-brain average of ≈5.5 reported meta-analytically (Blazey, et al., 2018), corresponding to a non-oxidative fraction of roughly 15–23% of glucose utilization. Visual stimulation lowered OGI by a further 12% in task-engaged regions — about two-fifths of the ≈30% reduction implied by the classic observations of Fox et al. (1988) — consistent with an activation-driven increase in aerobic glycolysis. Beyond the visual cortex, resting OGI was comparable (4.9) and did not differ systematically across functional networks, although individual frontal regions showed lower, and sensory regions higher, resting OGI than the cortical average. During stimulation, this wider cortex showed a smaller OGI reduction (≈8%) that, unlike in the visual cortex, was driven by concurrent decreases in CBF and CMRO₂ without significant changes in CMRglc or OEF. This reflects relative oxidative downregulation with preserved glucose utilization rather than true glycolytic upregulation, and should be confirmed with an independent measure of global OEF, for example, T2-relaxation-under-spin-tagging (TRUST) MRI (Lu & Ge, 2008). Overall, cortical metabolism is predominantly oxidative at rest but shifts toward glycolysis during stimulation — most strongly, and by a distinct mechanism, in the engaged visual cortex.

### Limitations

Several limitations should be considered. First, venous CBV was assumed to contribute 30% of total CBV changes, based on ultra-high-field data (J. J. Chen & Pike, 2010; Huber et al., 2014; Wesolowski et al., 2019), which may bias the CMRO₂ model (Fig. S2); direct measurement of venous CBV, for example, by combining LL-FAIR (Look-Locker flow-sensitive alternating inversion recovery) with contrast-agent techniques to separately quantify total and arterial CBV (Wesolowski et al., 2019), would reduce this source of uncertainty. Second, simultaneous mqBOLD–fPET was acquired with a 12-channel coil, limiting SNR and sensitivity to task-related perfusion changes; the relatively modest CBF (15-20%; Fig. 2) and comparable large total CBV increases (15-20%; Fig. S2) likely exerted opposing effects on CMRO₂ estimates, reducing detectable ΔCMRO₂; so higher-density coils and more sensitive perfusion sequences may yield larger and more accurate estimates of task-evoked oxygen metabolism. Third, baseline CMRO₂ values exceeded those obtained with the more reliable 3D T2-GRASE sequence (Epp et al., 2026; Raichle et al., 2001), probably because the 2D turbo spin-echo T2 mapping used here is susceptible to stimulated-echo contamination that inflates T2 and R2′; although corrections exist (Nöth et al., 2017), 3D T2-GRASE is now recommended (Kaczmarz et al., 2020), and future mqBOLD–fPET studies should adopt it or apply stimulated-echo correction. Because this bias primarily affects absolute CMRO₂, and hence resting OGI, rather than the within-session STIM–REST contrasts, it is unlikely to account for the task-induced OGI reductions reported here.

## Conclusion

This study shows that functional activation flexibly shifts cortical metabolism toward glycolysis. Simultaneous FDG-fPET and mqBOLD MRI revealed tightly coupled increases in CMRglc and CBF, only modest CMRO₂ changes, and clear OGI reductions in the visual cortex, alongside smaller, mechanistically distinct OGI reductions and CBF redistribution across the rest of the cortex. Beyond refining the interpretation of the BOLD signal as a spatially heterogeneous marker of neurovascular and neurometabolic coupling, the ability to track both energy substrates simultaneously opens new avenues for probing how altered vascular–metabolic interactions contribute to neurological and neuropsychiatric disease.

## Methods

### Participants

Twenty right-handed healthy participants were enrolled in the study. Four complete datasets were excluded due to excessive motion, SAR-related protocol modifications, or insufficient tracer administration. In addition, three mqBOLD datasets were excluded because of severe artifacts affecting at least one parameter map, and one fPET dataset was excluded as an outlier. The final sample included 15 participants with fPET data (29.9 ± 6.9 years; 9 females), 13 with mqBOLD data (33.1 ± 9.4 years; 8 females), and 12 with both modalities available (31.4 ± 7.0 years; 7 females). The study was approved by the Ethics Committee of the Klinikum rechts der Isar, Technical University of Munich (approval no. 334/13 S), and was conducted in accordance with the Declaration of Helsinki. All participants gave written informed consent prior to study participation.

#### Sample size

Using a two-sided significance level of 0.05 and a target statistical power of 95%, Reed et al. estimated that 13 participants were required for task-based [¹⁸F]FDG-fPET analyses using three-dimensional Gaussian spatial smoothing, comparable to the approach used here. We considered this estimate adequate for the present visual stimulation paradigm, as sensory tasks typically elicit stronger and more spatially focal activation than the higher-order cognitive tasks investigated by Reed et al. (2025).

### Experimental protocol

Participants arrived at the study site after an overnight fast. Baseline measurements (body weight, blood glucose, blood pressure, arterial oxygen saturation) were taken. An anesthesiologist placed two catheters: one intravenous in the left forearm for radiotracer and contrast agent, and one arterial in the right radial artery for blood sampling. Venous blood samples were collected for creatinine, hematocrit and glucose analyses. Participants were then moved into the scanner for imaging. During PET-MR scanning, participants experienced blocks of visual stimulation (STIM, a moving checkerboard at 8Hz) alternating with resting-state blocks (REST, a white fixation cross on a black background) approximately every 6 minutes, depending on the duration of the concurrent MR sequence.

PET scanning, visual presentation, and radiotracer infusion (3.6 MBq per kg body weight of 18F-FDG; Harvard Apparatus, Cambridge, Massachusetts, United States) started simultaneously. A 20% bolus injection (1ml/s) was administered initially to enhance signal-to-noise ratio (SNR) (Rischka et al., 2018), followed by a continuous infusion of the remaining 80% over 70 minutes (0.2ml/min). Continuous arterial blood sampling occurred throughout the PET duration using a Twilite system (Swisstrace, Zurich, Switzerland). Participants had to keep their eyes open during the experiment, as closed eyes significantly reduced metabolic activity in the visual cortex (Uludağ et al., 2004). A camera mounted above the head coil monitored participants’ wakefulness (MRC Systems GmbH, Heidelberg, Germany). Figure 1A shows a schematic depiction of the experimental protocol.

### Image acquisition

All data were acquired on a 3T Biograph PET-MR scanner (Siemens, Erlangen, Germany), using a 12-channel phase-array head-neck coil. Anatomical images were used for anatomical reference and to exclude brain lesions. This included a T1-weighted 3D MPRAGE pre- and post-gadolinium (TI=900 ms, TR=2300 ms, TE=2.98 ms, flip angle, α=9°; 160 slices, voxel size: 1.0×1.0×1.0 mm^3^; acquisition time: 5:03 minutes) and a T2-weighted 3D fluid-attenuated inversion recovery (FLAIR) image (TR = 5000 ms; TE = 394 ms, α=40°; 140 slices, voxel size: 0.5×0.5×1 mm^3^ EPI factor: 130, acquisition time: 3:27 minutes).

#### mqBOLD

The mqBOLD (Christen et al., 2012; Hirsch et al., 2014; Kaczmarz et al., 2020) protocol consisted of the following MR sequences:

- T2 mapping: 6:16 min 2D Turbo spin echo acquired only in REST (8 echoes, TE1 = ΔTE = 16 ms, TR=4870 ms, α=90°, voxel size 2×2×3 mm^3^, 36 slices).
- T2* mapping: 7:32 min multi-echo gradient-echo mapping acquired in STIM and REST (12 echoes, TE1 = 6ms, ΔTE = 5 ms, TR=2340 ms, α=30°, voxel size 2×2×3 mm^3^, gap 0.3 mm, 36 slices; 1 concatenation (4 concatenations in one subject), as described in (Hirsch et al., 2014; Kaczmarz et al., 2020). The images were corrected for magnetic background gradients with a standard exponential excitation pulse (Baudrexel et al., 2009; Hirsch & Preibisch, 2013), and half-resolution data of the k-space center were acquired for motion correction (Nöth et al., 2014).
- Arterial spin labeling: 5:09 min pseudo-continuous arterial spin labeling (pCASL) acquired in STIM and REST (post-labeling delay (PLD): 1800 ms, label duration: 1800 ms, 4 background suppression pulses, 2D EPI readout, TE=22.12 ms, TR=4600 ms, α=180°, 24 slices, EPI factor: 31, acquisition voxel size: 3×3×6.6 mm^3^, gap: 0.6 mm, 30 dynamic scans, including a proton density weighted M0 scan, following recommendations in (Alsop et al., 2015).
- Dynamic susceptibility contrast (DSC): 2.38-min single-shot GRE-EPI sequence (TE = 40 ms, TR = 1890 ms, α = 70°, voxel size = 2×2×3.5 mm³, 27 slices; 80 dynamics) acquired after a gadolinium-based contrast agent was administered as a bolus (0.2 mL/kg; two injections of 0.1 mL/kg for the two conditions, except for two subjects only in REST condition) after the first five dynamics (Hedderich et al., 2019), followed by 20 mL saline at 4 mL/s. Contrast administration was restricted to participants with normal renal function (creatinine ≤ 1.2 mg /dL).

#### fMRI BOLD

Additionally, we acquired 4:08min BOLD fMRI task block data using single-shot EPI (EPI factor: 64, voxel size = 3.0×3.0.3.0 mm^3^, FOV: 192×192×192mm^3^, TE=30 ms, TR=2.0 s, α=90°, 120 dynamic scans plus 2 dummy scans, 36 slices, interleaved acquisition together with a 0:54 min B0 field mapping scan (2 echoes, TR=400 ms, TE1=4.92 ms, TE2=7.38 ms, α=60°, voxel size: 3×3×3 mm^3^, 36 slices, interleaved acquisition).

### Data processing & statistical analyses

#### mqBOLD processing and CMRO_2_ calculation

Quantitative MR parameter maps were calculated with in-house scripts, using MATLAB and SPM12 (Wellcome Trust Centre for Human Neuroimaging, UCL, London, UK). Quantitative *T2* and *T2\** maps were obtained by mono-exponential fits of the multi-echo spin and gradient echo data (Hirsch et al., 2014; Kaczmarz et al., 2020; Preibisch et al., 2008). Corrections were performed for macroscopic magnetic background fields (Hirsch & Preibisch, 2013) and motion using redundant acquisitions of k-space center (Nöth et al., 2014). *R2’*, the transverse, reversible relaxation rate, was calculated via

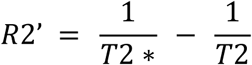

*R2’* depends on the vascular deoxygenated hemoglobin (dHb) content (Blockley et al., 2013b, 2015; Bright et al., 2019). Relative cerebral blood volume (CBV) was derived from DSC maps via full integration of leakage-corrected ΔR2*-curves (Boxermann, J.L., Schmainda, K.M., Weisskoff, R.M., 2006) and normalized to a white matter value of 2.5% (Hedderich et al., 2019; Kluge et al., 2016; Leenders et al., 1990). Combining *CBV* and *R2’* subsequently yielded the relative oxygen extraction fraction (OEF) via

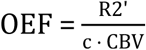

(Christen et al., 2012; Hirsch et al., 2014; Yablonskiy & Haacke, 1994), where c = γ · 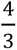· π · Δ χ0 ·hct · B0, the gyromagnetic ratio γ = 2.675 · 10^8^ s^-1^ T^-1^, the susceptibility difference between fully deoxygenated and oxygenated hemoglobin Δ χ_0_ = 0.264 · 10^-6^, the magnetic field strength B0 = 3T and the small-vessel hematocrit hct, which was estimated as 85% of subject-specific (large-vessel) hematocrit levels (Eichling et al., 1975; Hirsch et al., 2014). CBF maps were obtained from pCASL data as in Alsop et al. (2015) to calculate *CBF* from averaged, pairwise differences of motion-corrected label and control images and a proton-density weighted image. Finally, for each subject and condition, we calculated the voxelwise *CMRO_2_* by combining all parameter maps via Fick’s principle (Fick, 1870):

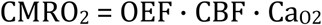

where *Ca_O2_* is the arterial oxygen content, calculated as *Ca_O2_* = 0.334 · *hct* · 55.6 · *O_2_sat*, with *O_2_sat* being the arterial oxygen saturation measured by pulse oximetry and *hct* the subject-specific hematocrit value (Bright et al., 2019; Ma et al., 2020). All parameter maps of each individual subject were registered to the first echo of their respective multi-echo T2 data. Figure 1C summarizes the calculation steps and shows the across-subjects averaged parameter maps in native space.

##### Case-specific adjustments

For two subjects, only a REST DSC acquisition was available. STIM CBV maps were therefore estimated by applying the mean task-evoked CBV change observed in the remaining subjects (n = 11) to each subject’s REST CBV.

##### Venous versus arterial CBV increases

CBV measured with dynamic susceptibility contrast (DSC) reflects total blood volume, including both arterial and venous compartments (Hua et al., 2019). While venous CBV changes can be neglected for brief stimuli (<40 s) due to their slow dynamics (Simon & Buxton, 2015), prolonged stimulation induces substantial venous CBV increases, accounting for approximately 30 (Huber et al., 2014) to 50% (Kim & Ogawa, 2012) of the total CBV response. Because visual stimulation lasted several minutes, CBV was measured in both conditions using separate DSC acquisitions. Since DSC provides total rather than venous CBV, group-level CMRO₂ estimates assumed that 30% of the observed CBV change was venous, based on human 7T visual stimulation data (Huber et al., 2014). A comparison of total versus venous CBV assumptions is shown in Supplementary Figure S2. As CBV was normalized to 2.5% and venous CBV was estimated rather than directly measured, both CBV and OEF should be interpreted as relative rather than absolute measures.

##### Artifacts and GM masking

To exclude susceptibility-affected regions, we computed voxel-wise temporal signal-to-noise ratio (tSNR) maps from BOLD-fMRI data in MNI space. Voxels within the lowest 15th percentile of tSNR in more than 66% of participants were excluded, primarily affecting frontobasal and temporobasal areas (Castrillon et al., 2023). The cerebellum was additionally removed, and analyses were restricted to voxels with gray-matter probability >0.5 to avoid unreliable OEF estimates in white matter due to oriented fibers. The resulting GM-SNR mask was applied to all partial least squares analyses.

R2′ is influenced not only by deoxyhemoglobin (dHb) but also by susceptibility-related confounds, including magnetic field inhomogeneities near air–tissue interfaces, iron accumulation in deep gray matter, myelinated white-matter fiber orientation, and tissue lipid content (Hirsch & Preibisch, 2013; Kaczmarz et al., 2020). Consequently, the native-space cortical (gray-matter) mask used in all analyses excluded voxels affected by cerebrospinal fluid (T2 > 150 ms), susceptibility artifacts (R2′ > 11 s⁻¹), large vessels (CBV > 12%), or physiologically implausible values (T2* > 90 ms, OEF > 1, CBF > 120). Absolute (Δ) and relative (%) task effects were computed as STIM − REST and (STIM − REST)/REST, respectively. OGI outliers were excluded using a robust criterion (OGI > median(OGI) + 3 × median absolute deviation) (Blazey, et al., 2018). Cortical OGI analyses were performed using the Schaefer 100-region parcellation (Schaefer et al., 2018), with each ROI assigned to one of the seven canonical Yeo functional networks (Thomas Yeo et al., 2011)(Fig. 3B).

#### PET processing, task analysis and CMRglc calculation

PET data were reconstructed offline from raw list-mode acquisitions using ordered-subsets expectation maximization (OSEM). Dynamic images were reconstructed into 93 frames of 45 s and a final frame of 15 s using a 3D iterative reconstruction (matrix = 344, zoom = 2.0, 4 iterations, 21 subsets, all-pass filter, relative scatter correction). Subsequently, the reconstructed PET images were motion-corrected using FSL mcflirt, temporally low-pass filtered (12-order Butterworth filter with 360 s cut-off) using scipy, and spatially iteratively smoothed in each direction (FWHM = 6mm) using AFNI 3dBlurToFWHM.

For CMRglc quantification, subject-specific arterial input function (AIF) was derived from continuous arterial blood sampling and processed using in-house Python scripts (v3.8). Blood-to-brain delay was estimated from the time between tracer injection and the arterial peak. The AIF was derived from blood time-activity curves (TACs) modeled from continuous arterial blood samples using a three-exponential function (Feng et al., 1993). The fitted whole-blood TACs were background-corrected and converted to plasma TACs using the FDG plasma-to-blood ratio model (Phelps et al., 1979) and subject-specific hematocrit values. The resulting plasma TAC, representing the AIF, was then evaluated at the midpoint of each reconstructed PET frame. For two participants with invalid arterial sampling, a population-based AIF was generated by averaging the remaining subjects’ AIFs (Castrillon et al., 2023; Vriens et al., 2009).

To account for the delayed measurable FDG uptake, task onsets were shifted by 2 min (Stiernman et al., 2021). fPET data were then analyzed using GLM with our Python implementation of the fPET toolbox (Hahn et al., 2026), modeling REST as the mean time series across non-occipital cortical gray matter, excluding task-expected activation regions, and STIM as a linear ramp with a unitary slope during stimulation periods and flat otherwise (Hahn et al., 2016). The STIM regressor was orthogonalized with respect to REST to reduce collinearity. REST and STIM tracer uptake trajectories were reconstructed from the corresponding GLM coefficients and combined with the processed AIF to estimate the net uptake rate constant (Ki) using Patlak analysis (Patlak & Blasberg, 1985). Finally, CMRglc was calculated as Ki multiplied by plasma glucose concentration and divided by a lumped constant of 0.65 (Wu, 2003).

#### fMRI BOLD processing and task analysis

The BOLD-fMRI localizer was preprocessed with fMRIPrep 20.2.4 (Esteban, 2019) running in a Docker container and based on Nipype 1.6.1 (Gorgolewski et al., 2011). Preprocessing included tissue segmentation, motion and confound estimation, susceptibility distortion correction, BOLD-to-T1w coregistration, and normalization to MNI152 ICBM 2009c Nonlinear Asymmetric space (2 mm). fMRIPrep uses FSL 5.0.9 (Jenkinson et al., 2012; Smith et al., 2004) tools BBR for BOLD-to-T1w registration, and FAST for tissue segmentation, and ANTs 2.3.3 for nonlinear spatial normalization (Avants et al., 2008), applying all transforms in a single interpolation step. The resulting normalization transform was also applied to all mqBOLD parameter maps following rigid (6-dof) coregistration to native T1w space for the GLM second-level analyses. The MNI Schaefer 100-region parcellation was transformed into mqBOLD space by concatenating the inverse mqBOLD-to-T1w and T1w-to-MNI transformations, enabling ROI-based analyses in native mqBOLD space.

First-level task analyses followed current recommendations (Esteban et al., 2020) and were conducted using a GLM including CSF and white-matter signals, DVARS, framewise displacement, and six motion parameters (translations and rotations) as confounds, together with a 120-s high-pass filter and 6-mm spatial smoothing.

#### Partial least squares analysis

mqBOLD maps task-evoked differences between REST and STIM were identified using mean-centered partial least squares (PLS) implemented in the Python pyls library. Unlike BOLD-fMRI and fPET, which provide multiple time points per condition for GLM analysis, mqBOLD parameters yield a single quantitative map per condition. Thus, PLS was used to identify the multivariate spatial patterns that maximally differentiated REST from STIM across subjects. PLS is a multivariate dimensionality-reduction approach that extracts latent variables (LVs) that capture the maximal covariance between brain signals and experimental conditions (McIntosh & Lobaugh, 2004; McIntosh & Mišić, 2013). The gray matter masked voxel-wise values from both REST and STIM conditions were entered directly into the analysis. Using a dummy-coding matrix, the pyls library computes within-condition averages and mean-centers them column-wise (Krishnan et al., 2011). Statistical significance of each LV was assessed using 2,000 permutation tests, and voxel reliability was estimated using 2,000 bootstrap resamples. Significant effects were identified using bootstrap ratios (BSR), computed as the ratio of voxel saliences to their bootstrap standard errors; |BSR| > 2 was considered significant, approximately corresponding to a 95% confidence interval under the assumption of normality (Krishnan et al., 2011; McIntosh & Mišić, 2013).

Group-level PLS mask corresponded to brain regions differentiating STIM from REST with a |BSR| > 2 and cluster size larger than 30 voxels. BSR maps were interpreted together with the corresponding design scores to determine the direction of the effect.

#### Other statistical analyses

The group-level GLM mask matched the second-level one-sample t-test (|Z-score| > 3.1) of the first-level activation maps across subjects. Hemodynamic and metabolic differences between REST and STIM were assessed using paired-sample t-tests across subjects. Cortical resting OGI values were evaluated using one-sample t-tests against the subject-averaged cortical OGI for each ROI, while task-evoked OGI changes (ΔOGI) were tested against zero across subjects. Multiple comparisons across ROIs were controlled using the Benjamini–Hochberg false discovery rate (FDR) procedure. Differences in ΔOGI across the 7-Yeo-networks were assessed using a one-way repeated-measures ANOVA, with *network* as a within-subject factor.

## Acknowledgments

We thank our colleagues at the Technical University of Munich, particularly C. Zimmer, W. Weber, G. Schneider, and S. Schachoff, who have consistently supported the multimodal imaging pipeline at TUM.

## Author contributions

Conceptualization: V.R. Supervision: V.R. and G.C. Data curation: S.E., A.B., G.C. and R.B. Investigation: S.E., A.B., R.B., K.K., E.C.D, A.R., E.S.B., M.B., I.Y., C.P., G.C. and V.R. Methodology: S.E, G.C. and R.B. Formal analysis: S.E., A.B., G.C., E.C.D., E.S.V and R.B. Visualization: S.E., A.B., G.C. and V.R. Validation: S.E. and G.C. Writing—original draft: S.E., A.B., G.C. and V.R. Writing—review and editing: S.E., A.B., R.B., K.K., E.C.D, A.R., E.S.B., M.B., I.Y., C.P., G.C. and V.R.

## Data and code availability

Images and derivatives are available in the OpenNeuro online repository (https://openneuro.org/datasets/ds008786), and the code is available on GitHub (https://github.com/NeuroenergeticsLab/fpet_mqbold_vistim).

