## Supplements for "Region-specific uncoupling of oxygen and glucose metabolism in the human cortex during visual stimulation"

Supplementary Materials

### Task-changes within group-level activation regions, for N=11 complete datasets

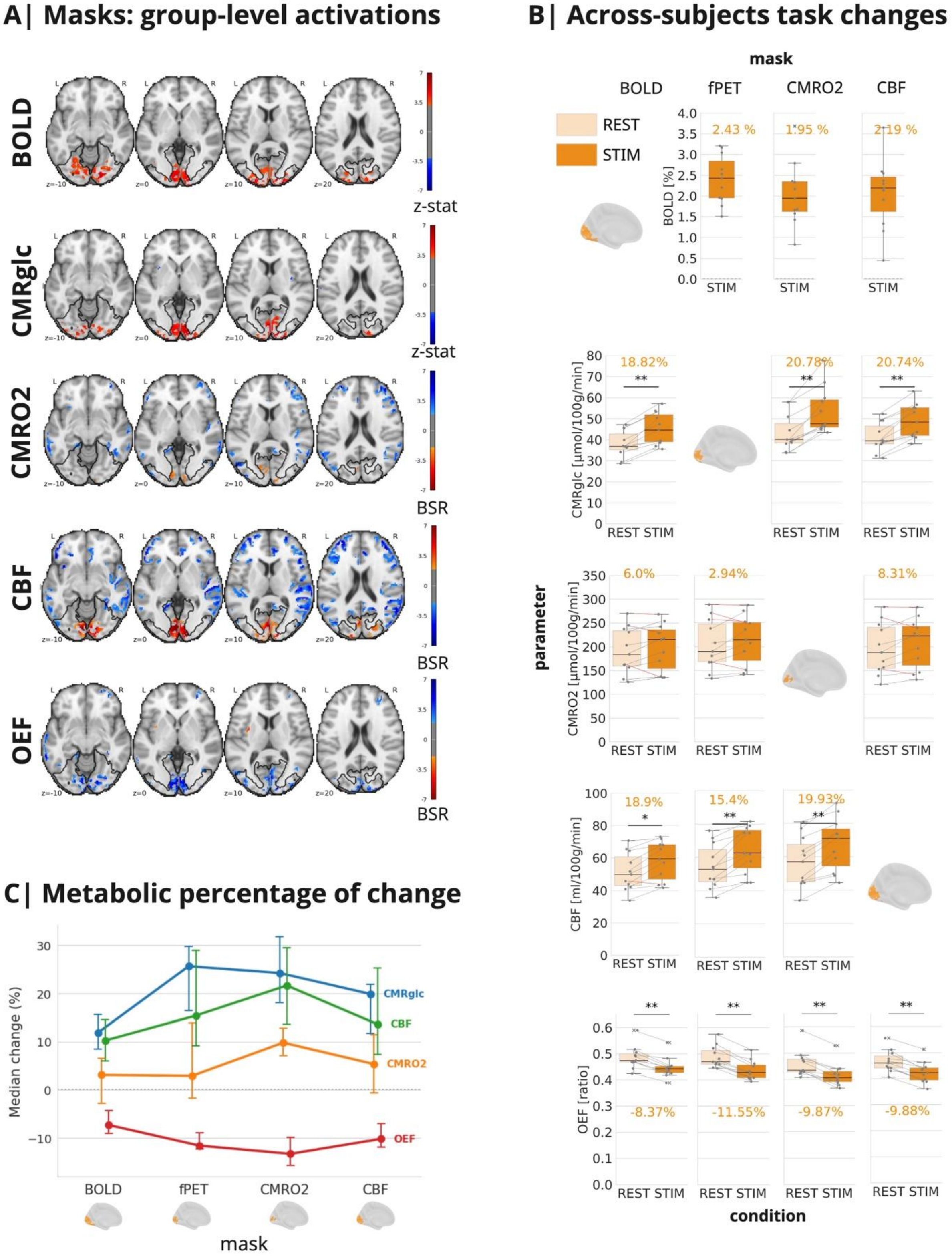

**Figure S1**. **Replication of task-evoked responses in the subset with complete DSC data**. Group-level maps and ROI-based analyses were repeated using only participants with DSC acquisitions in both REST and STIM conditions (N = 11). Results closely matched those from the full dataset (Fig. 2), confirming that the main findings were not driven by the two datasets without DSC during the STIM condition. **A**) Group- level STIM vs. REST maps in the subset with complete DSC data, showing increased BOLD, CBF, $CMRO_2$, and CMRglc, and decreased OEF within the visual cortex. **B**) Subject- level task- evoked changes within parameter- specific ROIs, all based on N=11 datasets. Results remained consistent with the full cohort, including the limited spatial specificity of $CMRO_2$ responses. **C**) Group-median percentage changes across ROIs. The relative magnitude of effects was preserved, with the largest changes in CMRglc and CBF, followed by OEF and $CMRO_2$. Error bars show 95% bootstrap confidence intervals.

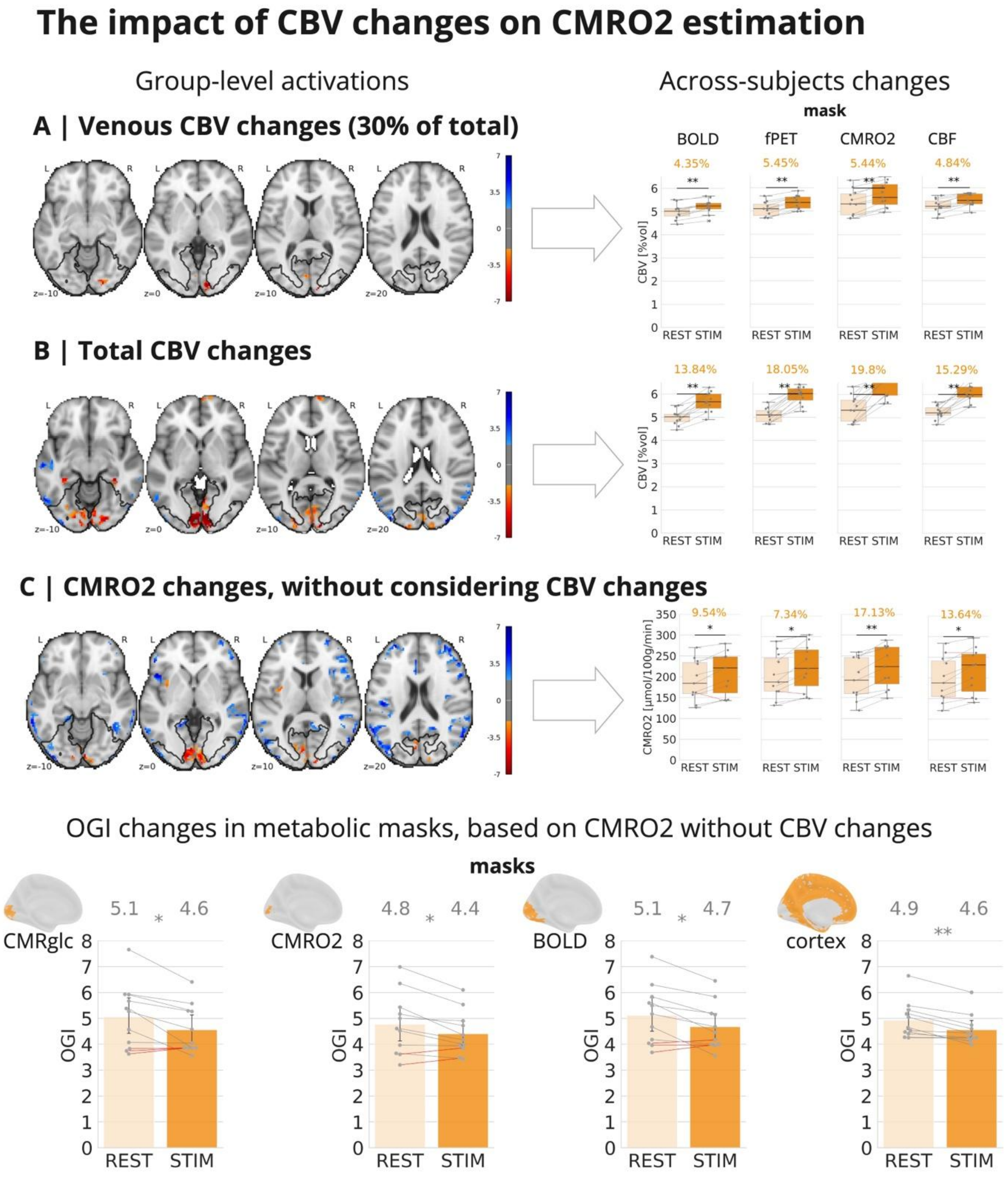


**Figure S2. Impact of CBV assumptions on $CMRO_2$ and OGI. A**) Group-level changes in venous CBV, corresponding to 30% of total CBV changes. $CMRO_2$ changes were calculated assuming venous CBV (main analysis). Left: group-level STIM vs. REST maps. Right: subject-level CBV changes within parameter-specific masks. **B**) Equivalent analyses using total CBV changes rather than estimated venous CBV changes. **C**) $CMRO_2$ changes calculated without accounting for task-evoked CBV changes. Upper-left: group-level maps. Upper-right: subject-level $CMRO_2$ changes across parameter-specific masks. Bottom: OGI changes within CMRglc-, $CMRO_2$-, and BOLD-defined ROIs, as well as the remaining cortex. Although omitting CBV correction increased estimated $CMRO_2$ responses, significant OGI reductions were preserved across all ROIs and the rest of the cortex, demonstrating that the main conclusions are robust to CBV assumptions.
